# Distinct gene expression profiles in leukocortical demyelinated white and grey matter areas of Multiple Sclerosis patients

**DOI:** 10.1101/2020.06.03.131300

**Authors:** T.A. van Wageningen, E. Gerrits, A. Geleijnse, N. Brouwer, J.J.G. Geurts, B.J.L. Eggen, H.W.G.M. Boddeke, A-M. van Dam

## Abstract

Demyelination of the CNS is a prominent pathological hallmark of Multiple Sclerosis (MS) and affects both white (WM) and grey matter (GM). However, demyelinated WM and GM areas exhibit clear pathological differences, most notably the presence or absence of inflammation and activated glial cells in WM and GM, respectively. In order to gain more insight into the differential pathology of demyelinated WM and GM areas, we micro-dissected neighbouring WM and GM demyelinated areas as well as normal appearing matter from leukocortical lesions of human post-mortem material and used these samples for RNA-sequencing. Our data show that even neighbouring WM and GM demyelinated areas share only 10% overlap in gene expression, implying a distinct gene expression profile, which is extending to a specific glial cell related signature. We propose that, based on their distinct expression profile, pathological processes in neighbouring WM and GM are likely different which could have implications for the efficacy of current MS treatments.

## INTRODUCTION

Multiple Sclerosis (MS) is a neurological disorder pathologically characterized by inflammation, demyelination and axonal damage in the central nervous system (CNS). It is the most common immune-mediated neurological disease among young adults, clinically represented by heterogeneous symptoms including sensory and motor deficits as well as fatigue and cognitive dysfunction^1^. Although demyelination is the pathological hallmark of MS, large differences exist between demyelinated white matter lesions (WMLs) and grey matter lesions (GMLs). Post-mortem analyses reveal that demyelination in the white matter is accompanied by activation of local glial cells and infiltration of peripheral leukocytes including monocytes and lymphocytes. In contrast, demyelinated grey matter areas show a paucity of activated glial cells and little to no infiltration of peripheral leukocytes^2, 3, 4^. Several hypotheses have been put forward as to why WMLs and GMLs differ in their pathology including (1) the presence of neuronal cell bodies, suppressing an immune response in the GM^5, 6^, (2) a difference in the abundance of myelin eliciting an immune response ^7^ or (3) that GM pathology is largely driven by the presence of meningeal infiltration of B-cells instead of parenchymal infiltration of leukocytes as is observed in the WM^8^. In addition, microglia present in normal appearing WM (NAWM) and normal appearing GM (NAGM) CNS tissue of MS donors show a specific transcriptional profile^9^, suggesting that regional differences in microglia may influence their response during MS pathology.

Analyses of human MS and related animal model brain material, suggest that, irrespective of the presence of infiltrating leukocytes, glial cells may be regulated during the pathological process in a region-specific manner^10, 11 4^. Chronic activation of microglial cells has generally been considered to play a detrimental role in WMLs whereas their role in GMLs remains unclear^2, 4, 11, 12^. Astrocyte scarring, represented by hypertrophic astrocytes and deposition of extracellular matrix proteins, can be identified most clearly in WMLs and is considered to impair regeneration^13, 14, 15^. Recent development in RNA single cell sequencing techniques have elucidated that under pathological conditions, e.g. Alzheimer’s disease and Multiple Sclerosis, subgroups of microglial cells can be identified based on their transcriptome which can have specific roles in the pathological process^16, 17^. Whether the transcriptomes of glial cells in WMLs and GMLs during MS are different is unexplored, and may reveal novel insight into their role in WM and GM pathology.

Here we addressed whether 1) genes are differentially expressed in GMLs versus WMLs, compared to corresponding normal appearing matter tissue and 2) if glial cell-related genes are differentially expressed in GMLs versus WMLs. Histologically verified human post-mortem leukocortical lesions were subjected to laser-capture microdissection (LCM) followed by RNA seq, to generate transcriptomic data of WML and GML areas within the same lesion and to compare these to corresponding NAWM and NAGM. Using leukocortical lesions that contained WM and GM demyelination within the same lesion minimizes possible confounding factors that could affect the results such as lesion location and time of lesion development. Our study showed that within a leukocortical lesion, demyelinated WM and GM areas shared less than 10% of genes that are changed in their expression compared to the corresponding normal appearing matter. Gene regulation in glial cells varied between demyelinated WM and GM areas with more pronounced regulation of astrocytic genes in WMLs, and of microglial genes in GMLs. Thus, distinct gene expression profiles in demyelinated WM and GM areas of leukocortical lesions may imply that different processes may underlie WML and GML pathogenesis which may open avenues for novel treatment targets implicated in WM and/or GM pathology.

## RESULTS

### Lesion characterization before laser capture microscopy

The presence of leukocortical lesions was assessed using consecutive frozen sections stained with antibodies against proteolipid protein (PLP) and MHC-II. Leukocortical lesions were defined by a loss of PLP immunoreactivity indicating demyelination in GM and WM (Fig. 1a). Leukocortical lesions are typically considered to be a type of GML and are therefore not scored based on the presence of MHC-II^+^ cells, however in order to facilitate interpretation of the subsequent RNA sequencing results from these lesions, we scored the WM demyelinated area of the leukocortical lesion based on MHC-II+ as is the convention for WMLs^18, 19^. Based on the presence of (1) MHC-II+ cells observed in a rim surrounding the hypocellular WM demyelinated area (Fig. 1b-d), (2) axonal transection as present by an increase in APP immunoreactivity (Fig. 1e) and (3) the presence of infiltrated CD3^+^ T-cells (Supplementary Fig. 1a,b), we scored the WM demyelinated areas of the leukocortical lesions as chronic active. The GM demyelinated area of the leukocortical lesions did not show an increase in MHC-II+ cells compared to the surrounding normal appearing matter. To help localize the lesion in the sections used for LCM, the sections were stained using a 4% cresyl violet which does not affect RNA-integrity^17^ (Fig. 1f-h). For readability, WM and GM demyelinated areas of the leukocortical lesion are referred to as WML and GML and normal appearing WM and GM as NAWM and NAGM respectively.

**Figure 1:**
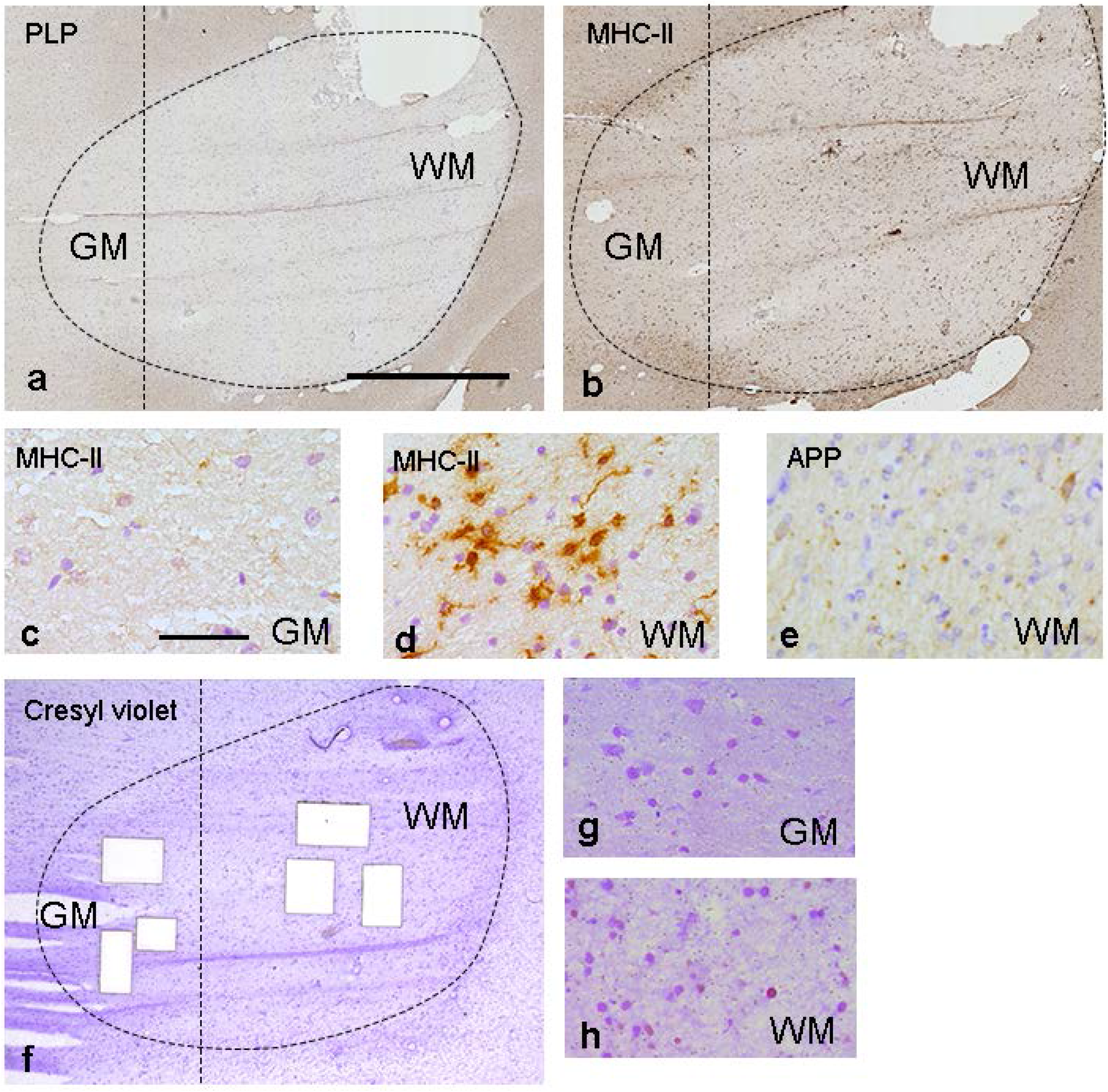
Immunohistochemical representation of the pathology of leukocortical lesions. (a) Loss of myelin as shown by reduced PLP staining in GM and WM area of the lesion, (b) Overview of MHC-II immunoreactivity in a leukocortical lesion, (c) Little MHCII immunoreactivity in GML, (d) Numerous MHC-II^+^ cells in WML, even though both areas are part of the same leukocortical lesion, (e) Presence of APP immunoreactivity in WML, indicative of axonal transection, (f) Tissue sections used for LCM were stained with 4% cresyl violet to localize (g) GM and (h) WM areas of the leukocortical lesion. Per section, 3 squares of tissue (white squares) per area were captured and pooled (f). Scalebar =500μm (a,b,f) or 25 μm (c-e,g,h).

### Quality control of RNA isolated from laser captured micro-dissected tissue

Tissue blocks from 11 MS patients (see Table 1) containing leukocortical lesions were included in the LCM and RNA seq analysis. Samples of 8 of these patients were subsequently used for gene expression data analysis (see Supplementary Data 1). In total 6 WML and 8 GML, 8 NAWM and 8 NAGM RNA seq samples were included in further analyses. To verify that LCM dissected tissue from different regions as input for RNA sequencing yielded representative data, we used principal component analysis (PCA). We observed that WM and GM samples segregated in the first principal component (PC1) (Fig. 2a). In addition, WML and NAWM clearly segregated in PC2, whereas GML and NAGM did not clearly segregate (Fig. 2a). This suggests that gene expression differences were more pronounced between WML and NAWM than between GML and NAGM. We identified 2352 genes which were differentially expressed (FDR < 0.05) between the 4 areas (GML, NAGM, WML, NAWM) of the leukocortical lesions. A heatmap of the top 50 genes with the lowest FDR (Fig. 2b) showed that the 4 areas of leukocortical lesions differentiated mostly in their expression of myelin (e.g. *PLP1, MAG*) and oligodendrocyte related genes (e.g. *OPALIN, CNP*) which were highest expressed in the NAWM followed by NAGM and as expected showed low expression in WMLs and GMLs. In addition, a clear separation was found in the expression of neuronal related genes (e.g. *NEFL, NEFH, GABRG2*) which was higher in GM areas than WM areas. Taken together, our data indicates that we were able to reliably dissect areas representing NAWM, NAGM, WMLs and GMLs from cresyl violet stained leukocortical lesions.

**Table 1:**
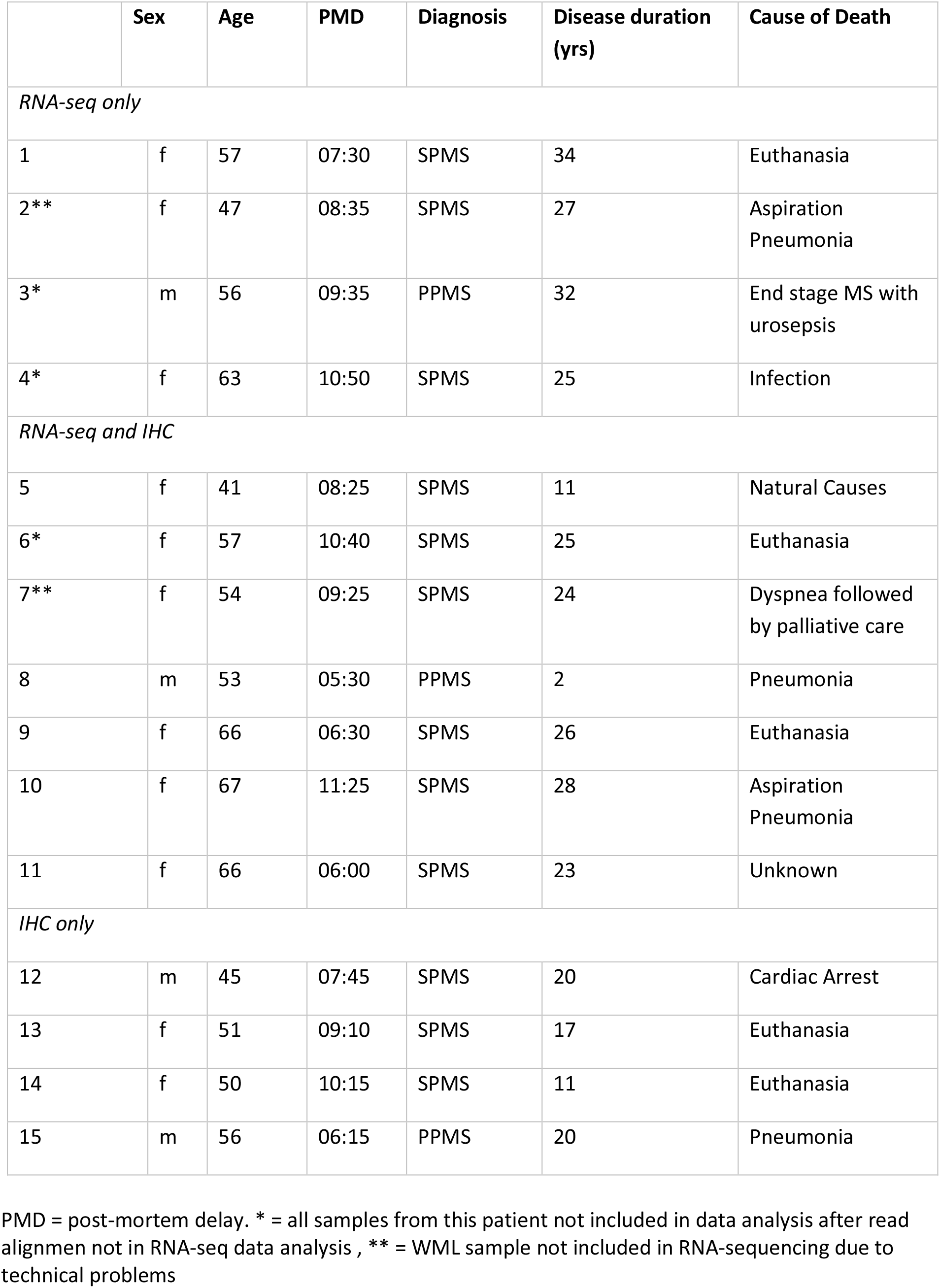
Clinical data of patients of tissue used for RNA-seq and immunohistochemistry

**Figure 2:**
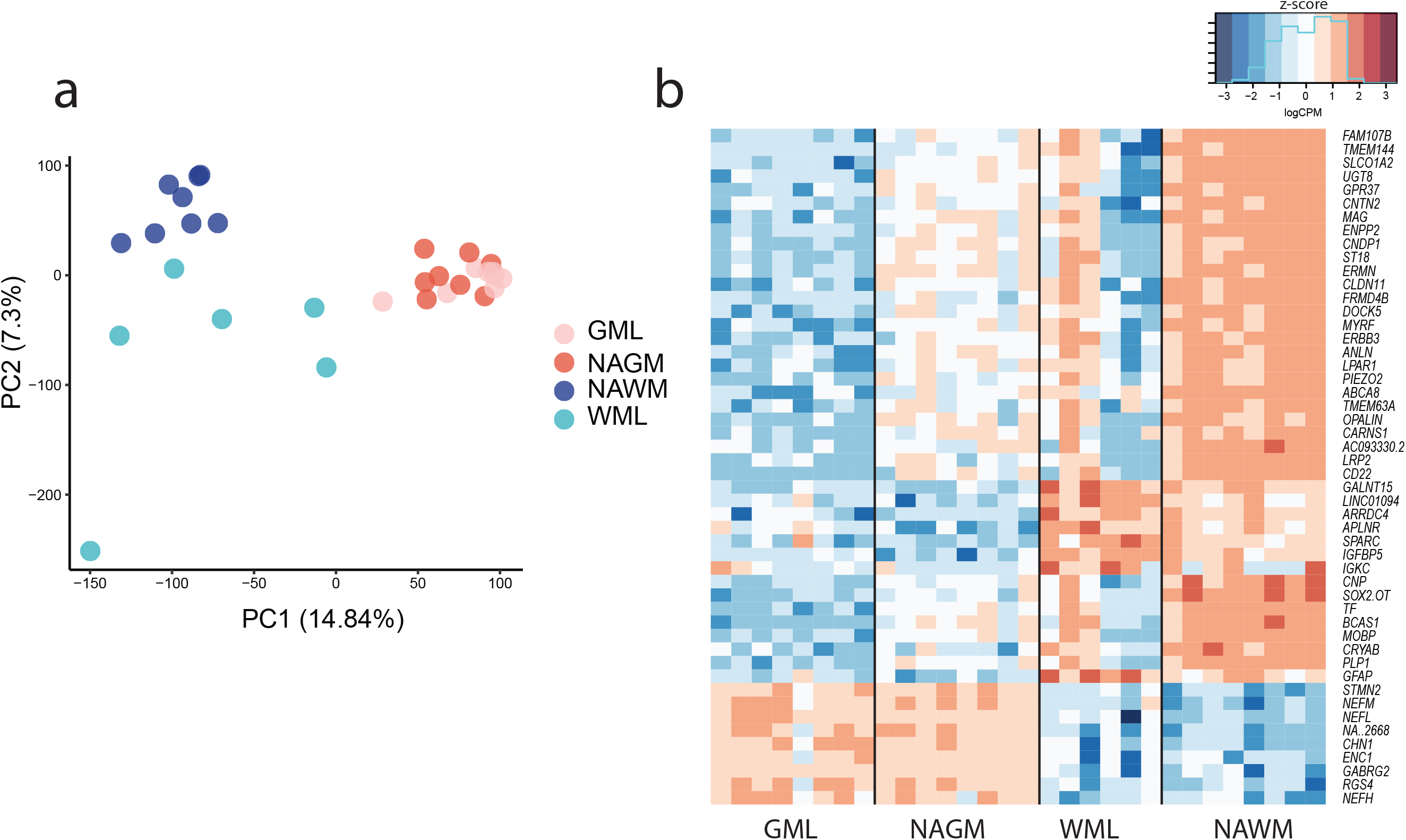
Overall differences in gene expression between NAWM, NAGM, WML and GML. (a) PCA plot of all samples included in RNA-sequencing showing separation of WM and GM derived samples as well as WML and NAWM derived samples. Separation of GML from NAGM samples is less clear, (b) Heatmap (log CPMs) of the top 50 genes differentiating between WML, GML, NAWM and NAGM samples according to the LRT-test. N= 8 for NAGM, NAWM and GML samples, N=6 for WML samples

### WMLs and GMLs within leukocortical MS lesions have distinct gene expression profiles

As can be observed from the heatmap (Fig. 2b), large differences in gene expression were present in NAWM compared to NAGM. Therefore, in order to gain insight into specific expression gene expression patterns of WMLs and GMLs, we contrasted gene expression of WML samples with NAWM samples and GML samples with NAGM samples. In total 2339 genes were significantly differentially expressed (logFC > (−)1, p <0.05) between WMLs and NAWM of which 760 were higher expressed in WMLs (Fig. 3a) and 1579 showed higher expression in NAWM (Fig. 3b). The comparison of GMLs vs. NAGM yielded a lower amount of 721 differentially expressed genes of which 462 were higher expressed in GMLs (Fig. 3a) and 259 were higher expressed in NAGM (Fig. 3b). As we showed that WMLs and GMLs exhibit different pathological characteristics, we wondered if this was also reflected in their gene expression. Indeed, only 10% (N=171) of the regulated genes overlapped between WMLs and GMLs, i.e. showed significantly higher or lower expression in WMLs and GMLs compared to NAM (grey areas Venn diagrams Fig. 3a,b). Not surprisingly, most of the shared genes lower expressed in WMLs and GMLs (N=141) were myelin or oligodendrocyte related, such as *MOG* and *CNDP1* (Fig. 3d). Shared genes that were higher expressed in both WMLs and GMLs were lower in number (N=30) and included genes related to inflammation such as *CD24* and *IGKC* (Fig. 3c). To determine whether the differential gene expression profiles of WMLs and GMLs may point to different biological processes, we conducted a pathway analysis on sets of genes which were higher expressed in WMLs or GMLs only (see Fig. 3e,f). Pathway analysis of genes expressed higher in GMLs compared to NAGM indicated that in GMLs there was increased regulation of cell adhesion and extracellular matrix organization related genes (Fig. 3g). Pathway analysis of genes expressed higher in WMLs compared to NAWM revealed that in WMLs there was increased regulation of genes related to cell-cell signaling but interestingly also synaptic signaling (Fig. 3h).

**Figure 3:**
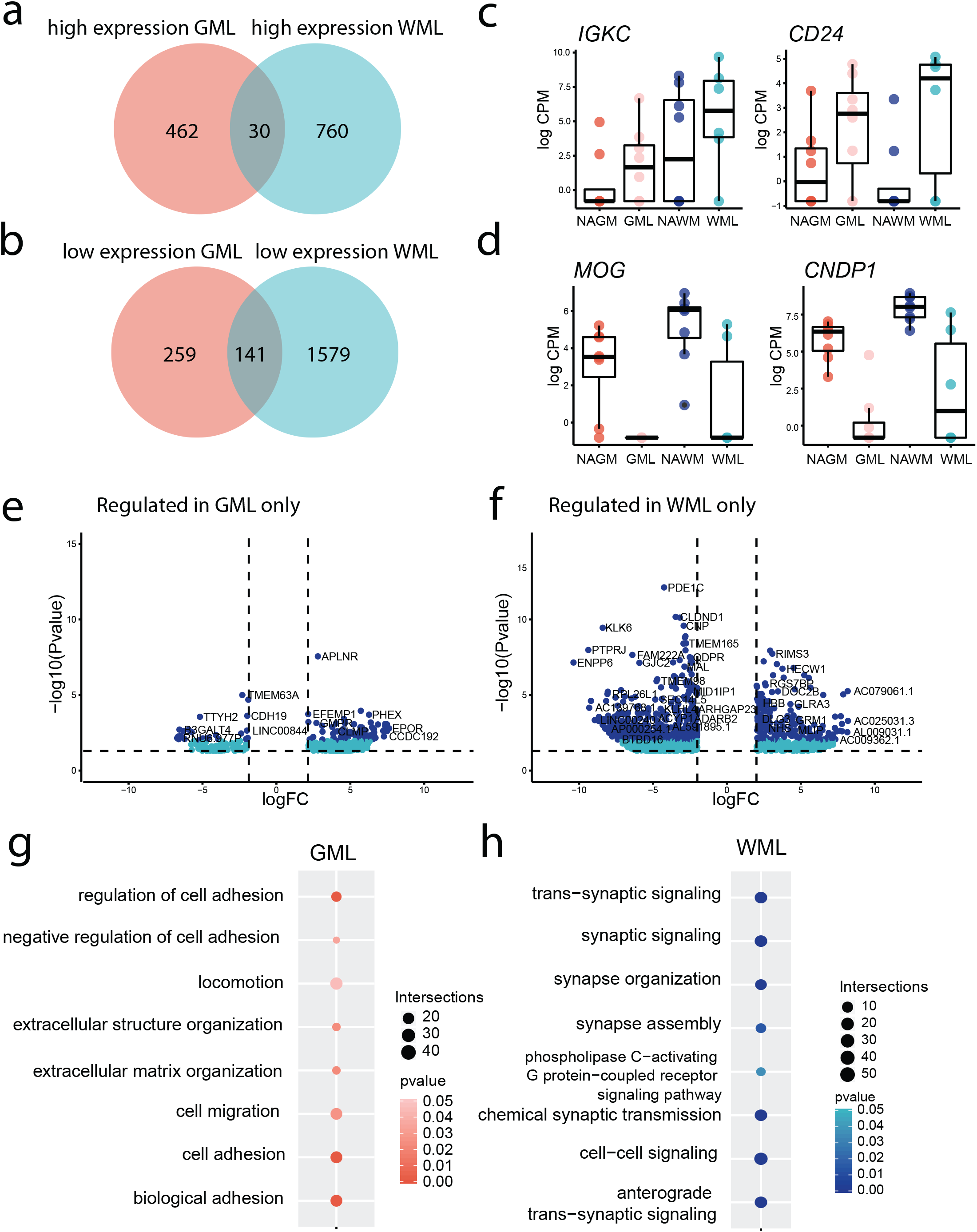
Limited overlap in gene expression between WMLs and GMLs. (a) Venn diagram showing the amount of genes higher expressed (logFC >2, p-value <0.05) specific to and overlapping between GMLs and WMLs compared to their respective NAM, (b) Venn diagram showing the amount of genes lower expressed (Log <−2, p-value <0.05) specific to and overlapping between in GMLs and WMLs compared to their respective NAM, (c) Graphs of genes higher expressed in both WMLs and GMLs compared to their respective NAM, (d) Graphs of genes lower expressed in both WMLs and GMLs compared to their respective NAM, (e) Volcano plot of genes differentially expressed (logFC <(−) 2, p value <0.05) in GMLs but not in WMLs, (f) Volcano plot of genes differentially expressed (logFC <(−) 2, p value <0.05) in WMLs but not in GMLs. Dark blue dots indicate genes logFC>(−)2 and p value <0.01, (g) Plot showing the pathways of genes enriched in GMLs but not in WMLs, (h) Plot showing the pathways of genes enriched in WMLs but not in GMLs. Intersections = number of genes represented in that pathway.

Tables of the differentially expressed genes in NAGM vs. NAWM, GML vs. NAGM and WML vs. NAWM, in addition to gene lists of overlapping and opposing genes regulated in WMLs and GMLs, can be found in Supplementary Data 2. In addition, pathway analysis for NAGM vs. NAWM, GML vs. NAGM and WML vs. NAWM can be found in Supplementary Data 3.

### Glial related changes in gene expression in WMLs and GMLs are not driven by changes in cell number or reactivity in the lesion

As recent transcriptomic data identified distinct glia subpopulations even within the healthy brain^9, 20, 21, 22^, we were particularly interested in the differences in gene expression between WMLs and GMLs by endogenous brain cells, e.g. local glial cells and neurons/axons. We therefore compared our gene lists of differentially expressed genes (log FC > (−) 1, p-value <0.05) in lesions and NAM with gene lists reported by Zhang et al.^23^, obtained by RNA sequencing of purified human brain cell types. We observed that in GMLs compared to NAGM, microglial and astrocyte related genes were most prominently higher expressed among the differentially expressed endogenous brain cell related genes (Fig. 4a), whereas oligodendrocyte related genes dominated the lower expressed genes (Fig. 4b). Interestingly, in particular microglial associated genes were higher expressed in GMLs suggesting a potential involvement of these cells also in seemingly low-inflammatory GMLs^2,24^. Compared to NAWM, in WMLs mostly neuronal/axonal and astrocyte related genes were higher expressed among the differentially expressed endogenous brain cell related genes (Fig. 4c), whereas, similar to GMLs, oligodendrocyte related genes were most abundant among the lower expressed in WMLs (Fig. 4d).

**Figure 4:**
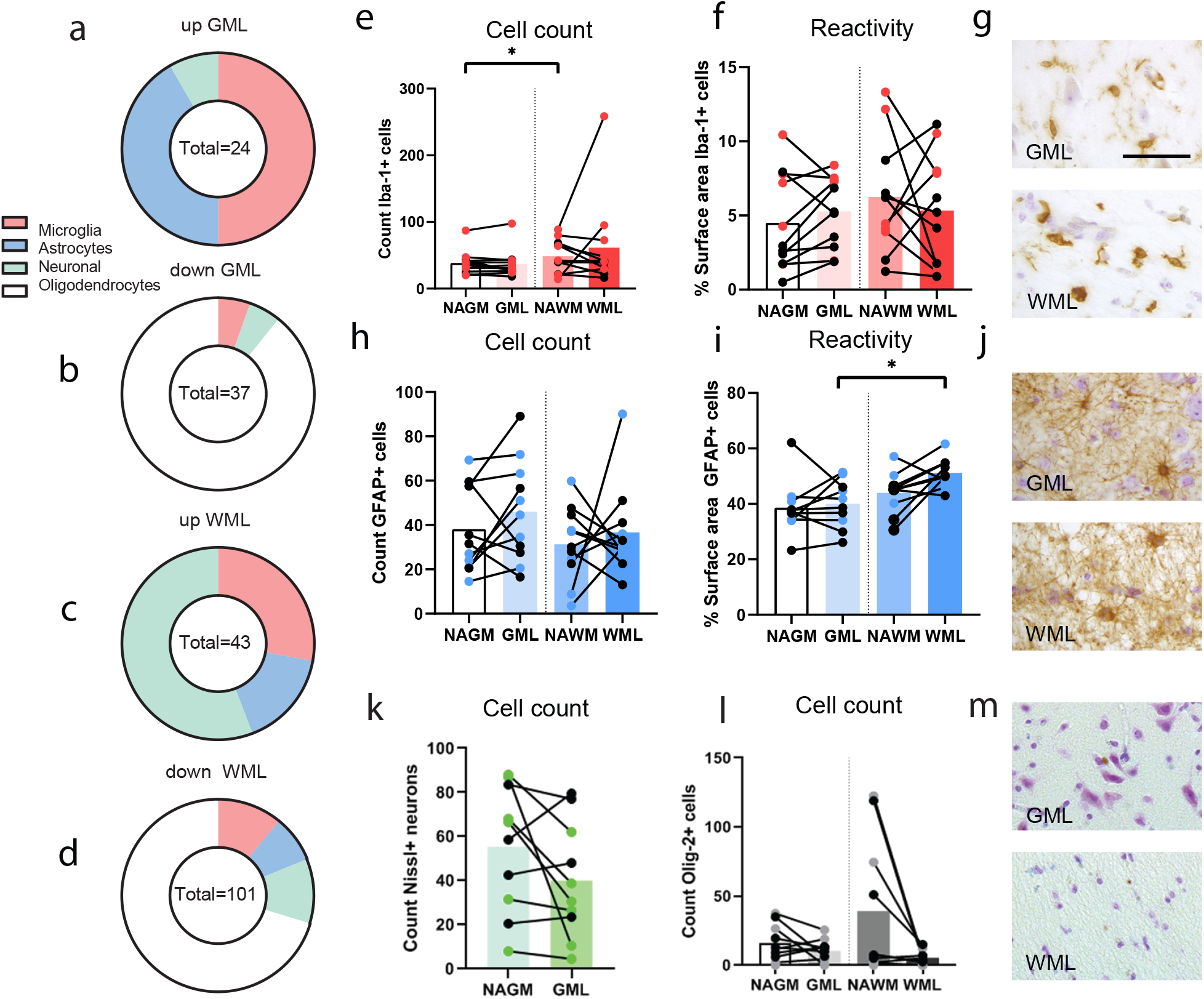
Overview of differentially regulated glial genes in WMLs and GMLs coupled with immunohistochemical analysis of standard glial cell markers to evaluate glial cell count and reactivity. Number of microglia, astrocytic, neuronal and oligodendrocyte (endogenous brain cells) related genes showing (a) increased expression (logFC >2, p-value <0.05) in GMLs compared to NAGM and (b) decreased expression (logFC <−2, p-value <0.05) in GMLs compared to NAGM, (c) Number of endogenous brain cell related genes increased (logFC >2, p-value <0.05) in WMLs compared to NAWM and (d) decreased (logFC <−2, p-value <0.05) in WMLs compared to NAWM. Graphs representing the amount (e) and reactivity (f) of Iba-1^+^ microglia in lesions compared to NAM, (g) Representative images of Iba-1^+^ microglial cells in GML and WML. Graphs representing the amount (h) and reactivity (i) of GFAP^+^ astrocytes in lesions compared to NAM, (j) Representative images of GFAP^+^ astrocytes in GML and WML, (k) Graph representing the cell count of Nissl^+^ neurons in NAGM compared to GML, (l) Graph representing the cell count of OLIG2^+^ oligodendrocytes in lesions compared to NAM, (m) Representative images of OLIG2^+^ oligodendrocytes in GMLs and WMLs. Coloured dots represent data obtained from same patients included in RNA-seq (e,f; h,I; k,l). Scalebar = 50 μm (g,j,m); *p<0.05.

To determine whether the observed difference in gene expression was due to actual gene regulation and not to a region-dependent loss or increase of specific cell types, we studied expression of cell type specific markers such as IBA-1 (microglia), GFAP (astrocytes), OLIG2 (oligodendrocytes) and Nissl (to identify neuronal cell bodies) on formalin fixed paraffin embedded (FFPE) material from MS patients identical to those used for RNA sequencing and from additional MS patients (see Table 1). No difference in microglia IBA-1 immunoreactivity was observed (F(3,30) = 1.834, p=0.162, Fig. 4f,g). Moreover, the number of IBA-1^+^ cells was not different between the four tissue areas (F(3,30)=3.047, p = 0.09, Greenhouse-Geisser corrected), but more IBA-1^+^ cells were counted in NAWM compared to NAGM (p<0.027) (Fig. 4e). The number of GFAP^+^ astrocytes was not different between the tissue areas (F(3,30)=1.131, p=0.354). However, GFAP^+^ astrocytes did show a group difference in reactivity (F(3,27)=5.581, p=0.004). This effect was mostly driven by the larger GFAP^+^ cell surface areas in WMLs compared to GMLs (p = 0.034)(Fig. 4h-j). Loss of Nissl^+^ neurons was not observed in GMLs compared to NAGM (t(9) = 1,751, p = 0.1138, Fig. 4k). Surprisingly, there was no statistical difference in the number of OLIG2^+^ cells when comparing all four tissue areas (F(3,21) = 2.329, p=0.15, Greenhouse-Geisser corrected, Fig. 4l), even though a decrease in OLIG2^+^ cells was visualized in lesioned areas (Fig. 4m). This is likely explained by the large variation in OLIG-2^+^ cell numbers within NAWM (Fig. 4l). We therefore eliminated oligodendrocyte related genes from further analysis to prevent loss of oligodendrocytes as confounding factor influencing data interpretation. In contrast, we observed no large differences in microglia, astrocyte or neuron numbers or reactivity in lesioned compared to normal appearing matter based on the immunoreactivity of the selected protein markers. Thus, the differential expression of cell related gene expression were considered to be due to actual regulation of gene expression.

### Differential gene expression of microglial related genes in WMLs and GMLs within leukocortical lesions

As both WML and GML samples of leukocortical lesions did not show an increase or decrease in IBA-1^+^ microglia, we explored next which microglial related genes did show increased expression in samples from WMLs and GMLs. Similarly to IBA-1, we found that both WMLs and GMLs did not show a no significant differences in expression of homeostatic microglia markers such as *TMEM119* and *P2RY12* (Fig. 5a,b), although a tendency to downregulation of *TMEM119* in the chronic active WM demyelinated area was present corresponding to what we reported before ^24^. In addition, we observed no difference in expression in leukocortical lesions of two known microglial activation markers for MS lesions, *CD74* and *HLA-DR* (Fig. 5a,b). Most genes found differentially expressed in GMLs compared to NAGM (Fig. 5c) showed no significant differential expression in WMLs compared to NAWM. Similarly, genes found significantly differentially expressed in WMLs were not significantly differentially expressed in GMLs (Fig. 5d). This suggests that the microglia population in WMLs and GMLs differs. Of interest is that known anti-inflammatory related genes *CD163* and *MRC1* were significantly higher expressed in WMLs only (Fig. 5d). Both *CD163* and *MRC1* are associated with a response to IL-4^25^. Other, less studied genes in the context of MS showing increased expression in WMLs such as e.g. *TRIM38* and *KLF10* are involved in suppression of immune responses^26, 27^. These data suggest that significantly higher expressed microglia related genes in WMLs are indicative of a dampened local immune response in leukocortical lesions.

**Figure 5:**
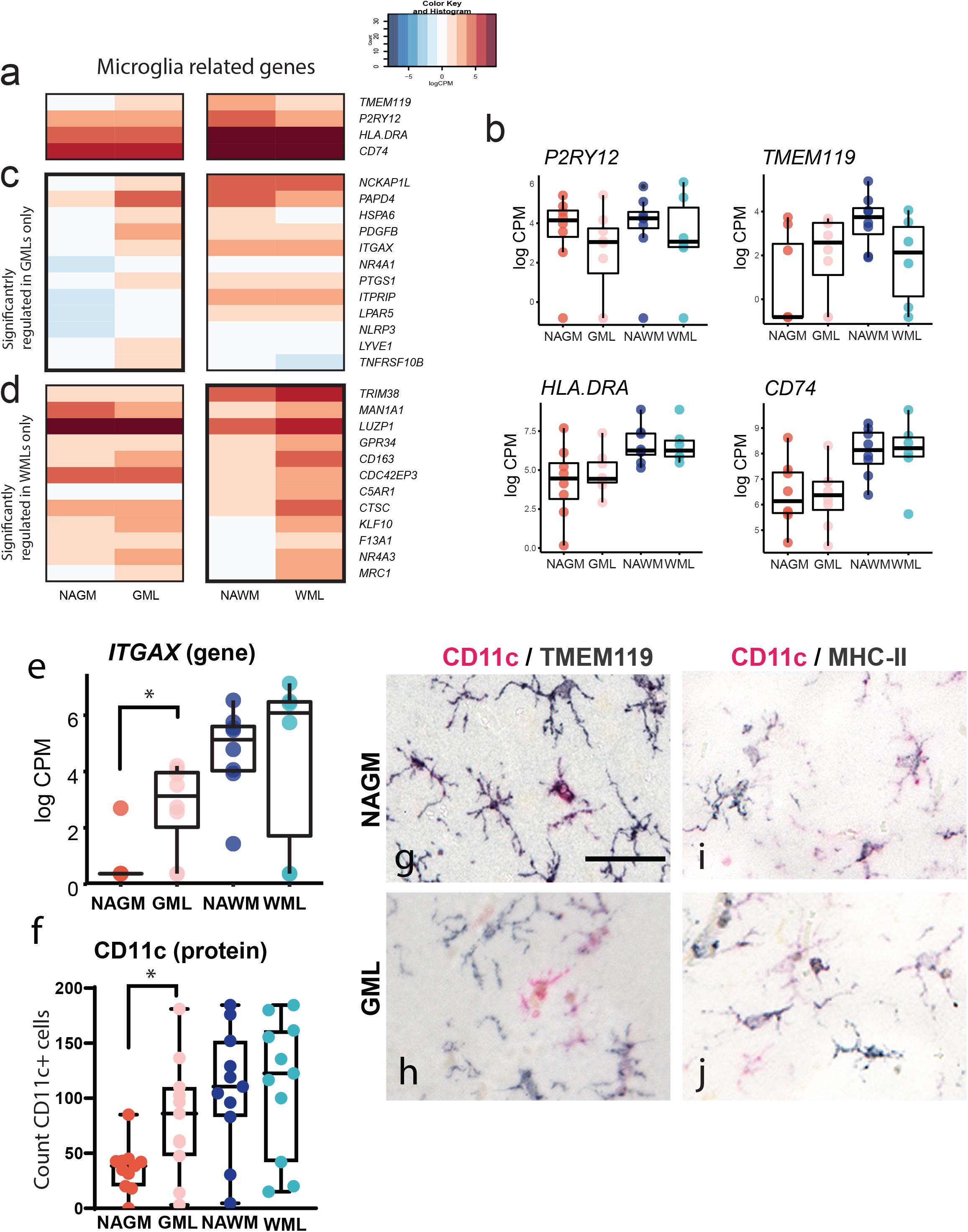
Overview of gene expression and protein validation of microglial related gene expression in WMLs and GMLs. (a) Heatmap of averaged logCPMs and (b) Graphs indicating no differential expression of known MS microglia markers such as *HLA-DR* and *CD74* as well as the homeostatic microglia markers *P2RY12* and *TMEM119*, (c) Heatmap of averaged logCPMs of NAGM, GML, NAWM and WML samples of significantly higher expressed microglial related genes (logFC >2, p-value <0.05) in GMLs only, (d) Heatmap of averaged logCPMs of NAGM, GML, NAWM and WML samples of significantly higher expressed microglial related genes (logFC >2, p-value <0.05) in WMLs only,(e) *ITGAX* expression determined by RNAseq, (f) Semi-automatic quantification shows an increase in number of CD11c^+^ cells in GMLs compared to NAGM only, (g,h) Representative images of CD11c andTMEM119, (I,j) CD11c and MHC-II in NAGM and GMLs. Scalebar = 100 μm (g-j). * = p<0.05

When studying the microglia related genes in GMLs that are significantly higher expressed than in NAGM, we observed that some of these genes e.g. *NLRP3, ITGAX* and *NR4A1* (Fig. 5c) have been reported to interact with the pro-inflammatory cytokine IL-1β^28,29,30^, which was also found higher expressed in GMLs (data not shown), suggestive for an inflammatory microglia profile that is however different from that in WMLs. Interestingly, GMLs also featured an upregulation of *PDFGB* and *LYVE1* which showed higher expression in GMLs than NAWM and NAGM (Fig. 5c), both of which are involved in chemotaxis but also in communication with the endothelium^16,31^. We subsequently aimed at validating a gene that was upregulated in GMLs only, opening opportunities to finding a GML specific microglia marker. To this end, *ITGAX* (Fig. 5e) was chosen for validation by immunohistochemistry. The number of CD11c^+^ (ITGAX) cells (F(3,27)=2.194, p=0.112) showed no significant difference when comparing all four tissue areas, however after post-hoc testing, a significant increase in the number of CD11c^+^ cells was found in GMLs compared to NAGM only (p=0.041, Fig. 5f). Using double-labeling immunohistochemistry, we found that CD11c^+^ cells also expressed TMEM119, supporting that indeed microglia express CD11c in both NAGM (Fig. 5g) and demyelinated GM (Fig. 5h). In addition, most, but not all, MHCII^+^ microglia showed co-localization with CD11c (Fig. 5i,j). These data highlight a subset of CD11c^+^ microglia that is increased in GMLs only.

### Differential gene expression of astrocyte related genes in WMLs and GMls within leukocortical lesions

Similar to microglia, astrocyte related upregulated genes did not show overlap between WMLs and GMLs of leukocortical lesions which may suggest different functions for astrocytes in WMLs and GMLs (Fig. 6a-c). Known astrocyte markers, often used to study astrocyte status in MS lesions, *GFAP* and *AQP4* showed significantly higher expression in WMLs compared to NAWM and no difference in expression between GMLs and NAGM (Fig. 6b). *SLC1A2*, which is known to be more selective for GM astrocytes ^32^ indeed shows higher gene expression in GM areas, but shows not regulation in WMLs or GMLs whereas *SLC1A3*, which is also proposed to be GM selective, shows no difference in gene expression between WM and GM and only a moderate increase in WMLs and GMLs^32^. (Fig 6b). Genes higher expressed in demyelinated GM only included e.g. *SDC2, GLI3, DOK5* and *CLDN10* (Fig. 6c) which also implicate a role for astrocytes in supporting neuronal function^33,34,35, 36^.

**Figure 6:**
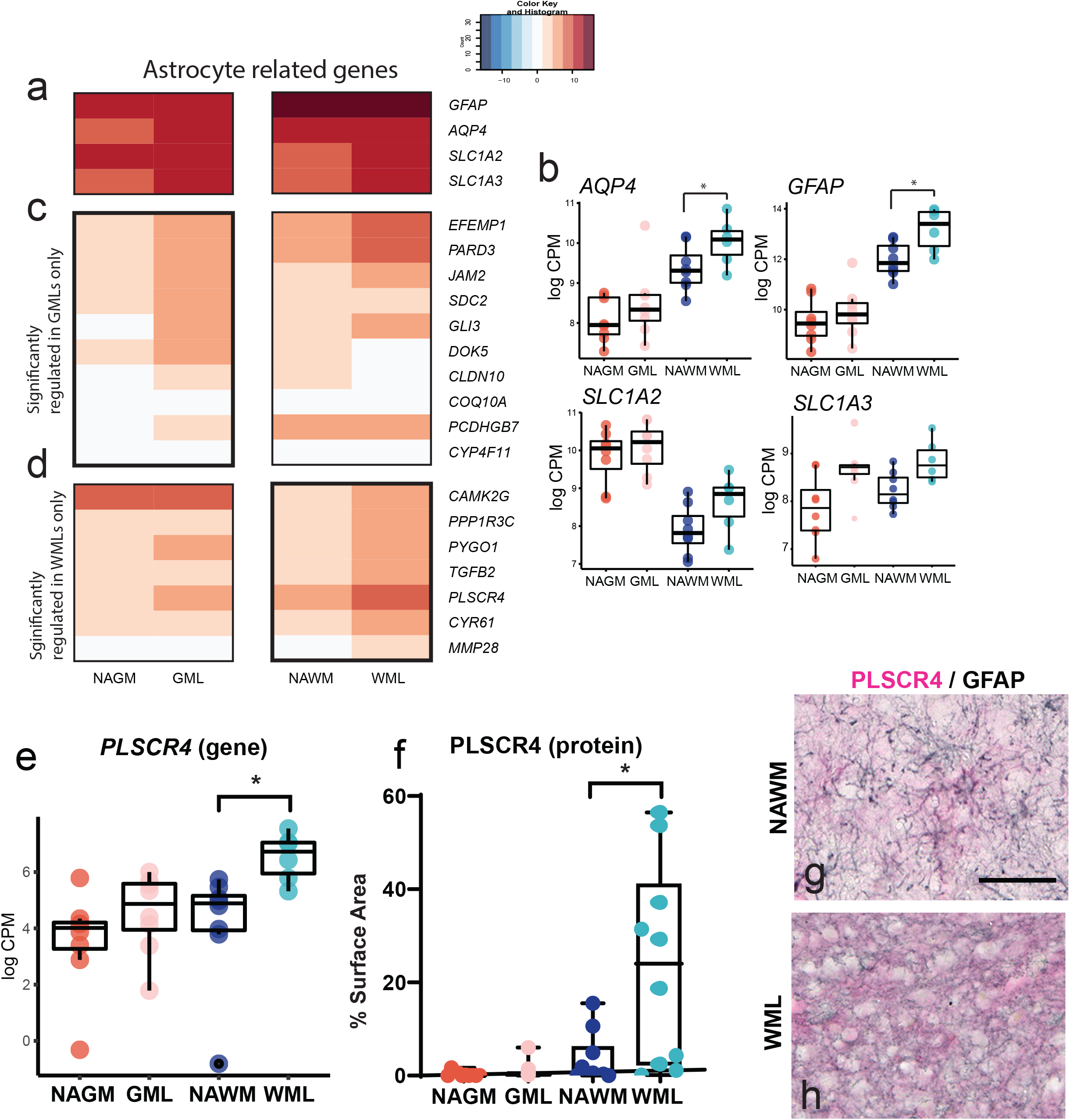
Overview of gene expression and protein validation of astrocyte related gene expression in WMLs and GMLs. (a) Heatmap of averaged logCPMs and (b) Graphs indicating regulation of known astrocyte genes *AQP4, GFAP, SCL1A2 and SCL1A3* in WMLs but not in GMLs, (c) Heatmap of averaged logCPM values of NAGM, GML, NAWM and WML samples of significantly higher expressed astrocyte related genes (logFC >2, p-value <0.05) in GMLs only, (d) Heatmap of averaged logCPM values of NAGM, GML, NAWM and WML samples of significantly higher expressed astrocyte related genes (logFC >2, p-value <0.05) in WMLs only, (e) *PLSCR4* expression as determined by RNAseq, (f) Semi-automatic quantification shows an increase in PLSCR4 immunoreactivity in WMLs compared to NAWM only, (g-h) Representative images of PLSCR4 and GFAP immunoreactivity in (g) NAWM and (h) WMLs. Scalebar = 100 μm (g,h). * = p<0.05.

Of genes regulated only in WMLs (Fig. 6d), *TGFB2* was higher expressed. *TGFB2* is considered to be a hub gene for chronic-active WMLs ^37^ but not in GMLs, highlighting again the differences in glial responses between WMLs and GMLs. Other, less explored, genes in the context of MS showing higher expression in WMLs included. *CYR61, CAMK2G* and *PLSCR4* (Fig. 6d). We chose to validate the expression of *PLSCR4* as it was most pronounced increased in WMLs (Fig. 6d,e). Using immunohistochemistry a significant group difference in the immunoreactivity of PLSCR4 (F(3,27)=17.817, p<0.000) was observed which was apparent for WMLs compared to NAWM (p=0.02, Fig. 6f). Double-labeling immunohistochemistry showed that PLSCR4 immunoreactivity was present in GFAP^+^ astrocytes in WMLs and NAWM (Fig. 6g,h), supporting an astrocytic source of PLSCR4. The presence of enhanced PLSCR4 immunoreactivity in astrocytes in WMLs only, opens avenues for PLSCR4 as marker for, at least a subset, of astrocytes in WMLs.

## DISCUSSION

Although white and grey matter pathology develops during the course of MS, grey matter pathology is more prominent in later stages of the disease^38,39^. Since GMLs are less inflammatory (e.g. there is a relative absence of infiltrating leukocytes), but do show extensive demyelination, we hypothesized that endogenous brain cells could contribute to either grey or white matter pathology in a significant manner. In the present study we aimed at identifying (glial) genes that are differentially expressed in WML and GMLs compared to their respective NAM. In order to minimize the effect of lesion location, we utilized LCM on snap-frozen sections to micro-dissect WMLs, GMLs and NAWM and NAGM from leukocortical lesions of MS patients featuring demyelination in neighboring WM and GM. This approach allowed us to micro-dissect areas from histologically verified leukocortical lesions allowing accurate localization of GM demyelination. However, as LCM can have a significant impact on RNA integrity, we chose to use bulk RNA seq to get accurate gene expression profiles. We chose to focus on leukocortical lesions as these lesions are defined by demyelination in neighboring WM and GM and thus we hypothesized that we minimized the interference of lesion location and lesion time of development on gene expression. However, it must be noted that the exact pathological mechanisms of leukocortical lesion development are still unknown.

When studying genes specific for WMLs and GMLs, a surprising finding was the little overlap in genes regulated in the WMLs and GMLs of leukocortical lesions. Less than 10% of differentially expressed genes in WMLs and GMLs compared to NAM overlapped, suggesting differential pathological processes in WMLs and GMLs even when part of the same leukocortical lesion. Indeed, pathway analysis of genes differentially expressed in WMLs-only indicated gene expression differences related to cell-cell signaling and synaptic signaling suggesting there is a substantial regulation of genes related to neurons in WMLs. Notably, GMLs did not feature an enrichment of these pathways and instead genes higher expressed in GMLs were associated with cell adhesion and regulation of the extracellular matrix. These opposing results, together with the very limited overlap of gene expression between WMLs and GMLs indicate different pathological processes are apparent within the same leukocortical lesion. Furthermore, gene expression in both WMLs and GMLs did not show an enrichment for leukocyte or lymphocyte related pathways even though CD3^+^ T-cells were observed in WM demyelinated areas, confirming that the role of infiltrated leukocytes might be less relevant in leukocortical lesions obtained from secondary progressive MS patients. We therefore focused on elucidating gene expression changes of local endogenous brain cells such as microglia and astrocytes.

Using immunohistochemistry we confirmed that the observed changes in gene expression in WMLs and GMLs are not due to a clear loss or increase of specific cell types. Therefore, it is likely that changes in glial related gene expression are due to regulation of genes, rather than cell loss. In order to gain insight into gene expression differences related to astrocytes and microglia, we extracted genes highly expressed by, and specific to microglia and astrocyte from cell specific gene sets. Gene expression of microglial related genes in WMLs showed an increased expression of *CD163* and *MRC1*. Microglia and or macrophages characterized by expression of *CD163* and *MRC1* have been shown to be involved in oligodendrocyte differentiation, thereby possibly contributing to efficient remyelination as shown in lysolecithin induced lesions in mice^40^. The upregulation of *CD163* and *MRC1* therefore suggests that in chronic-active WMLs of leukocortical lesions, microglia contribute to creating an environment supporting remyelination. Although we cannot exclude that *CD163* and *MRC1* are expressed by infiltrated, foamy macrophages in WMLs^41^, our cohort featured relatively little IBA-1^+^ foamy macrophages in WMLs of leukocortical lesions, indicating our data likely represents local microglia exhibiting an anti-inflammatory phenotype. Furthermore, increased expression of *TRIM38*, an ubiquitin-protein ligase which inhibits TLR3 mediated IFN type 1 signaling^26^ and *KLF10*, induced by TGFβ, which suppressed microglial activation^27^ also support a less inflammatory profile.

In contrast, whereas we did not observe increased expression of inflammatory genes commonly used to stage activation of lesions in MS such as *HLA-DR* and *CD74*, microglia related genes in GMLs did reflect an inflammatory status of microglia. We observed an increase in gene expression of *NLRP3* and *ITGAX.* Increased *NLRP3* expression can be induced by microglial mitochondrial damage, suggesting that microglia in GMLs are impaired in their mitochondrial functioning, something which is also observed in Parkinson’s disease^42^. In addition, activation of *NLRP3*, as part of the inflammasome, increases processing and release of the pro-inflammatory cytokine IL-1β which we also find increased in GMLs compared to NAGM. A role for IL-1β in MS pathophysiology has been clearly established^29,43^. Increased *ITGAX* expression in GMLs is also indicative of an activated microglial state: CD11c^+^ microglia have recently been described as an identifier for plaque related disease-associated microglia (DAM) in Alzheimer’s disease (AD) and Amyotrophic lateral sclerosis (ALS)^21^, thus pathology related microglia. Taken together, the increase in *NLPR3* and *ITGAX* in leukocortical GML microglia may represent a specific microglial subset in GMLs which is different from that observed in WMLs. By studying *ITGAX* at the immunohistochemical level (i.e. CD11c) we confirmed the selective regulation of CD11c in the GM area of leukocortical lesions and can thus be used as a potential new marker for a subset of microglia involved in the pathology of MS GMLs where there is no increase in *HLA-DR* expression at the gene and protein level^24^.

Similar to microglia, astrocyte-related regulated genes differ between WMLs and GMLs of leukocortical lesions. In our study, we observed selective regulation of the phospholipid scramblase *PLSCR4* ^44^ in demyelinated WM at the gene and the protein level. PLSCR4 can interact with CD4 which is expressed by T- helper- cells^44^. Recently, it was shown that the astrocytic response in the EAE animal model of MS is shaped by GM-CSF released by infiltrated T-helper cells in WMLs^45^. Thus, increased expression of PLSCR4 could reflect an increased communication between astrocytes and T helper-cells still present in chronic active WMLs. Possibly, the increased expression of *PLSCR4* is indicative of the role of astrocytes in modulating differentiation of CD4^+^ T-cells to T-helper (Th) 1 cells^46^. Though MS has long been considered a CD4+/Th-1 mediated disease^47^, this has been questioned and the presence of CD4^+^ Th-1 cells possibly induced by astrocytes could be indicative of neuroprotective events which is also reflected in the presence of *CD163* and *MRC1* expressed by microglia as previously discussed^48^. Indeed, astrocytes have also been shown to limit T-cell proliferation, including CD4^+^ T-cells, thereby mitigating CNS autoimmunity^46^. In contrast to WMLs, there was neither upregulation of *PLSCR4* nor of *GFAP, AQP4, SLC1A2* or *SLC1A3* in GMLs, indicating that the astrocyte response in leukocortical GMLs is rather different from that in WMLs. Instead, GMLs of leukocortical lesions featured upregulation of *SDC2* and *GLI3*, two astrocyte genes related of which the proteins syndecan-2 and zinc finger protein *GLI3* are involved in tripartite maturation through modulation of cell adhesion^33, 34^ and sonic hedgehog signaling^35^, respectively. The upregulation of *SDC2* and *GLI3* are suggestive of synaptic changes in GMLs. Astrocytes are crucial in removing glutamate from the synaptic cleft^49^. Activation of astrocytes by e.g. IL-1β (which was found increased in GMLs) leads to downregulation of glutamate transporters by astrocytes, preventing removal of glutamate from the synaptic cleft and leading to excitotoxic synaptopathy and thus synaptic loss^49^. Indeed synaptic loss has been found in MS hippocampus^50^ and during chronic demyelination of mouse brains^34^. Increased expression of *SDC2* and *GLI3* could suggest active regulation of the tripartite synapse by astrocytes in GMLs, possibly hindering further (neuronal) damage.

Taken together, our study highlights distinct gene expression profiles in demyelinated WM and GM areas in MS, within the same leukocortical lesion. Moreover, we observed differential regulation of glial cell-related genes in WMLS and GMLs. Highlighting that WMLs and GMLs are characterized by a different glial signature. Of interest is that numerous regulated glial genes have not been previously described in the context of MS and could give new insights into specific glial genes involved in either WM or GM pathology, to be addressed in future studies. Moreover, the limited overlap in (glial) related genes can have potential consequences for the development and efficacy of treatment for MS. Currently, treatment efficacy is monitored mostly based on inflammatory WMLs, as these are visible on conventional MRI due to high inflammatory activity likely induced by infiltrated leukocytes. However, the striking differences in (glial) gene expression of WMLs and GMLs, even when spatially close together indicate different pathologies ongoing in WMLs and GMLs. Further elucidating the cells and processes underlying WM and GM pathology in MS could lead to new treatment targets which might be more beneficial for treatment of progressive MS, when GM pathology becomes more prominent.

## MATERIALS AND METHODS

### Human brain tissue

Fresh-frozen tissue blocks and formalin-fixed paraffin embedded human brain tissue blocks were obtained from the Netherlands Brain Bank and the Amsterdam Multiple Sclerosis Centre, Amsterdam UMC. In compliance with all local ethical and legal guidelines, informed consent for brain autopsy and the use of brain tissue and clinical information for scientific research was given by either the donor or the next of kin. Clinical information can be found in table 1.

### Immunohistochemistry

Fresh frozen and formalin fixed paraffin embedded (FFPE) tissue blocks were sectioned at 10 μm and mounted on positively charged glass slides (Menzel-gläser, ThermoFisher Scientific). Cryo-section were fixed in 100% acetone for 45 min, dried and washed in 0.1 M Tris-buffered saline (TBS; pH 7.6) whereas FFPE sections were deparaffinized before undergoing antigen retrieval using 10 mM Tris buffer containing 1 mM EDTA (Tris-EDTA, pH 9). Subsequently, all sections were incubated for 20 min with 1% H_2_O_2_ in TBS to block endogenous peroxidase, washed in TBS and incubated for 30 min in 5% donkey serum in TBS with 0.1% Triton-X (block buffer) to block non-specific antibody binding. Thereafter, sections were incubated with primary antibodies (Suppl. Table 2) diluted in block buffer overnight at 4°C. Then, sections were washed with TBS and incubated in block buffer containing biotinylated donkey anti mouse IgG (1:400, Jackson laboratories) at room temperature (RT) for 2 hr. Subsequently, sections were washed in TBS and incubated for 1 hr with horseradish peroxidase-labeled avidin-biotin complex (ABC complex, 1:400, Vector Labs) at RT. For CD11c staining, Envision + HRP labelled polymer anti-rabbit (DAKO) was used as a replacement of the secondary antibody to enhance the signal. Immunoreactivity was then visualized by adding 3,3-diaminobenzidine (DAB, Sigma, St. Louis, USA) and sections were counterstained with hematoxylin. Sections were subsequently dehydrated in graded series of ethanol, cleared in xylene and mounted with Entellan (Merck).

Double-labeling immunohistochemistry was performed on FFPE sections treated similarly as described above until the incubation with primary antibodies. Sections were incubated with combinations of primary antibodies (CD11c + TMEM119 or CD11c + MHC-II or PLSCR4 + GFAP) diluted in block buffer, overnight at 4 °C (Suppl. Table 1). Sections were washed in TBS and incubated with ImmPRESS anti-Rabbit IgG polymer detection kit (Vectorlabs) for 30 min at RT. Subsequently, slides were washed again and incubated for 30 min at RT with Envision+ HRP labelled polymer anti-mouse (DAKO). Subsequently, immunoreactivity was visualized by adding Liquid Permanent Red (LPR, DAKO) to visualize CD11c or PLSCR4 and Vector SD Peroxidase kit (Vectorlab) to visualize TMEM119, MHC-II or GFAP. Subsequently, sections were washed in TBS and demi-water before being dried on a heat plate at 37°C before and cleared in xylene and mounted with Entellan (Merck).

### Laser capture microscopy

Sample preparation for laser capture microscopy (LCM) was done as previously published ^51^ with slight modifications. In short: tissue blocks were sectioned at 10 μm in an RNAse free cryostat and mounted on PEN-slides (Leica). Immediately after mounting, the sections were fixed in 100% molecular grade ethanol for 10 min and dried. Subsequently, the sections were stored at −80°C, paired back to back in a 50 ml tube containing silica gel as a desiccant. Upon use for LCM, sections were taken from storage and immediately put into 100% molecular grade ethanol, rehydrated to 70% ethanol and then stained in 4% cresyl violet in 70% ethanol for 45 sec. After staining, sections were dehydrated in an ethanol series (up to 100% ethanol). Sections were dried and then stored again in 50 ml tubes containing silica gel on dry ice before commencing LCM. Upon use for LCM, frozen, cresyl-violet stained sections on PEN-slides were transferred to 100% ethanol for 1-2 minutes and dried. NAGM, NAWM, GML and WML were localized and dissected using the Leica LMD6500 system (Leica, Germany). Per section, 3 squares of tissue per area were captured (Fig. 1c,f,g). Per patient, tissue was captured from 6-12 cresyl violet stained sections.

### RNA isolation and RIN determination

Pooled laser captured samples per area per patient were collected in 50 μl extraction buffer of the ARCTURUS^®^ PicoPure^®^ RNA Isolation Kit (ThermoFisher Scientific), incubated at 42°C for 30 min (according to the manufacturers protocol) and stored at −80°C until further use. RNA was isolated according to the manufacturers protocol with an additional DNA-se digestion step using the Qiagen RNAse free DNAse kit (Qiagen, Germany). RNA quality and concentration of all samples were determined using the Agilent Bioanalyzer before and after the LCM procedure.

In total, frozen tissue blocks from 11 MS patients featured a type I leukocortical lesion that fitted the previously established pathological criteria (see Fig. 1a,b and d, e). The mean ± standard deviation (SD) RNA integrity indicated by the RIN score of the tissue blocks before the LCM procedure was 6.2 ± 2. After the LCM procedure the mean ±SD RIN value of laser-captured samples was 5.1 ± 1.2 which was considered appropriate to be used for RNA sequencing.

### RNA sequencing

Sequencing libraries were prepared with the Quant Seq 3’ mRNA seq Library Prep Kit FWD (Lexogen, Vienna, Austria). The libraries were sequenced on a NextSeq platform.

### Data Analysis

Quality control of the raw FASTQ files was performed with FASTQC. Bad quality bases were trimmed with TrimGalore version 0.4.5. Sequences were aligned using HiSat2 version 2.1 to the *H. Sapiens* (GRCh38.92) reference template obtained from Ensembl and quantified with featureCounts. A quality check of aligned data was performed with FASTQC and MultiQC. Raw count matrices were loaded in R and annotated by converting the ensemble IDs to gene symbols using the corresponding gtf file. Only genes with > 1 counts in at least 2 samples were included in the analysis.

Subsequently, data was normalized and analyzed using the *EdgeR* package. A likelihood ratio test(LRT)-test was used to compare gene expression between the four defined groups using only lesion type as a variable (i.e. NAWM, NAGM, WML and GML). Subsequently an LRT-test design blocking patient variation (~0+lesiontype + patients) was used to produce gene lists contrasting GML vs. NAGM, WML vs. NAWM and NAWM vs. NAGM. Using these lists, sub-lists were created contrasting upregulated or downregulated genes in GML compared WML. Up- or down-regulated genes were identified with the criteria of a logFC >(−)2 and a P value of <0.05. Gene-set enrichment analysis of up- and down-regulated genes was conducted using the *g:Profiler* package. Differentially expressed (DE) genes were defined by a FDR <0.05 and logFC >(−)1. Other packages used were: *DEseq2* (for log2 transformation of the counts), *DEGpatterns* (to visualize the results of the group comparison), *ggplot2*, and *GeneOverlap.*

To determine the cell type specificity of genes, we used data generated by Zhang et al. (2016)^23^ utilizing RNA sequencing on purified astrocytes, microglia, oligodendrocytes and neurons derived from human brain. Data was downloaded from NCBI GEO, ascension number GSE3721. Using their FPKM normalized counts from human derived cells, we sorted the data such that we could extract the top 2000 genes highly expressed by each cell type (astrocytes, microglia, oligodendrocytes and neurons). We then compared these gene lists and subtracted genes that were also expressed by multiple cell types (e.g. oligodendrocyte gene OLIG2 was also expressed by astrocytes) to generate a proxy of highly expressed, genes per specific cell type. Total amount of cell-type specific genes in our dataset after this correction was microglia: N=715, astrocytes: N=392, oligodendrocytes: N=294 and neurons: N=499.

### Analysis immunoreactivity

Immunoreactivity detected in demyelinated WM and GM areas of leukocortical lesions and of corresponding NAWM and NAGM was analyzed using ImageJ. Per lesion, depending on the lesion size, 1-2 images were made at 20x magnification using a Leica DM5000B microscope. Per NAWM and NAGM area, 2 images were made. All images analyzed had a region of interest (ROI) of 622 × 466 μm. Within these ROIs, signals from DAB and hematoxylin were separated using the *color deconvolution* plug-in. From the subsequently acquired DAB images without hematoxylin signal, an auto-threshold method was applied and % area stained and cell counts were generated using the *Analyze Particles* plug-in. Neurons from Nissl stained slides were thresholded using an auto-threshold method and selected from all other cell bodies by setting in the *Analyze Particles* plug a predefined minimal size (>53 um^2^).

Cell counts for CD3 were conducted using an Olympus BX45 microscope with a U-OCMSQ 10/10 eyepiece micrometer (Olympus Lifescience) featuring a square of 10 mm2. Cells CD3^+^ were counted in three squares of 10 mm2 at 20x magnifications and counts were averaged and adjusted to a count of positive cells/mm2.

### Statistical analysis

Statistical analysis of quantitative data of immunohistochemical stainings was conducted using SPSS. Data of all stainings were not normally distributed and were therefore normalized using a log10 transformation. Since all four areas from one patient can be considered paired measurements, a repeated measures ANOVA was used with post-hoc testing using Bonferroni correction.

## Data availability

RNA sequencing data presented in this study have been deposited at https://www.ncbi.nlm.nih.gov/geo/ under the accession number: GSE149326 and will be made public upon acceptance for publication of this manuscript.

## Author Contributions

TvW performed and analyzed the experiments, interpreted the data and wrote the draft version of the manuscript; EG contributed to data analysis of RNA sequencing data, AG and NB contributed to experiment performance, JG, BE and HB contributed to the intellectual content and AvD conceived the study and contributed to the intellectual content. All authors contributed to and approved the final manuscript.

## Acknowledgements

We would like to acknowledge Kenn Zwaan and Frederike Dijk for technical assistance with respectively the RIN value determination and LCM.

This work was financially supported by the Dutch MS Research Foundation (grant no. 15-904MS received by A-M van Dam and H.W.G.M. Boddeke).

---

**Supplementary figure 1:** Graphs representing the number of CD3^+^ T-cells infiltrated in the (a) perivascular space and (b) parenchyma of leukocortical WMLs and GMLs compared to NAM. ** = p <0.01, **** = p<0.0001.

